# Unprecedented exoelectrogenic activity in *Parachlorella kessleri* MACC-38: The interplay of photosynthetic electron transport and the oxidative pentose phosphate pathway

**DOI:** 10.1101/2025.03.28.645967

**Authors:** Nia Z. Petrova, Soujanya Kuntam, Milán Szabó, Kashif Shaikh Mohd, Gyula Kajner, Gábor Galbács, Szilvia Z. Tóth

## Abstract

Exoelectrogenesis is the ability of living cells to export electrons. In photosynthetic organisms, exoelectrogenesis is of particular interest because it can be used for transduction of solar energy into electric current in biophotovoltaics or reducing power for biocatalysis. Previously, we identified and characterized the green microalga *Parachlorella kessleri* MACC-38 producing unprecedented current largely dependent on the photosynthetic electron transport chain (PETC). In this study, our comparative photosynthetic characterization demonstrates that chlororespiration is a crucial factor maintaining the PETC redox balance in MACC-38. Our data points that the oxidative pentose phosphate pathway (OPPP) is activated during exoelectrogenesis to meet the increased demand of reducing power. We hypothesize that chlororespitration prevents oversaturation of PETC during OPPP activation. These findings provide valuable insights into the fundamental mechanisms of exoelectrogenesis in green algae. By elucidating the complex interconnection between PETC, OPPP and exoelectrogenesis, our results pave the way towards improved bioelectrochemical and biocatalysis technologies.

## 1. Introduction

The ability of living cells to reduce compounds in their immediate environment (i.e., to export electrons) is known as exoelectrogenic activity. This phenomenon has been best studied in prokaryotic organisms owing to the availability of two model metal-reducing bacterial species *Shewanella oneidensis* MR-1 and *Geobacter sulfurreducens*, known as electricigens. In these organisms, exoelectrogenesis is an indispensable part of cell metabolism. Extracellular reduction of metal (hydr)oxides as terminal electron acceptors ensures respiratory activity in anaerobic conditions (Gao et al., 2010; Levar et al., 2014). Furthermore, the exoelectrogenic activity of these species exerts broad ecological impact, affecting the availability of growth-limiting or toxic compounds and organic matter in waters and soils, providing conditions for syntrophic interactions and more globally, ensuring the biogeochemical cycles (Summers et al., 2010; Williams et al., 2013).

Exoelectrogenesis has been observed at a significant scale in several photosynthetic microorganisms, including cyanobacteria, green microalgae, and diatoms (Xue et al., 1998; Weger and Espie, 2000; Laohavisit et al., 2015; Anderson et al., 2016; Gonzalez-Aravena et al., 2018; Petrova et al., 2024). Although the full range of physiological and ecological functions of exoelectrogenesis in algae is yet to be explored, it is known that electron export in these organisms facilitates extracellular reactive oxygen species formation (Anderson et al., 2011, 2016; Yuasa et al., 2020) and metal reduction (Hill et al., 1996; Xue et al., 1998; Gonzalez-Aravena et al., 2018). A key feature of microalgal exoelectrogenesis is its dependence on photosynthetic activity (Xue et al., 1998; Weger and Espie, 2000; Laohavisit et al., 2015; Yuasa et al., 2020). This dependence opens avenues for the transformation of sunlight into electric current in biophotovoltaic devices (BPV), also known as bio-photoelectrochemical cells, and into reducing power for the synthesis of value-added chemicals (Wey et al., 2021, 2024; McCormick et al., 2011; Anderson et al., 2016; Wang et al., 2021; Laohavisit et al., 2016; Vicente-Garcia et al., 2024; Petrova et al., 2024).

Our current understanding of the interplay between photosynthetic electron transfer reactions and electron export is largely based on studies employing common inhibitors. These studies have shown that blocking the photosynthetic electron transport chain (PETC) at photosystem II (PSII) or the terminal acceptors of photosystem I (PSI) inhibits electron export and bioelectricity production in cyanobacteria and green algae (Lynnes et al., 1998; Bombelli et al., 2011; Pisciotta et al., 2011; Gonzalez-Aravena et al., 2018; Shlosberg et al., 2021; Petrova et al., 2024). Further studies in cyanobacteria have revealed that the oxidative pentose phosphate pathway (OPPP) activity during the dark period preceding current production, along with respiration, provides reducing power for light-induced electricity generation (Saper et al., 2019; Hatano et al. 2022). Additionally, the deletion of the thylakoid-localized respiratory terminal oxidases has been shown to enhance electric current production in cyanobacteria (Bradley et al., 2013). In the green alga *C. reinhardtii,* NADPH has been identified as the primary direct electron donor for exoelectrogenic processes (Xue et al., 1998; Anderson et al., 2016; Wang et al., 2021). Under specific conditions, such as iron limitation, the NADPH used for electron export is derived from both the OPPP and the PETC (Xue et al., 1998). Overexpression of ferredoxin 1 or ferredoxin 5 in *C. reinhardtii* promotes current production, suggesting a role for ferredoxins as an electron cross-point between PETC, NADPH, and electrons destined for export (Huang et al., 2015). Treating green microalgal cells with electron acceptor potassium ferricyanide (FeCN) has been shown to augment photosynthetic oxygen evolution and alleviate photoinhibition, highlighting the role of exoelectrogenesis as an electron sink (Weger and Espie, 2000; Wang et al., 2021). Nevertheless, our understanding of the specific photosynthetic processes crucial for exoelectrogenesis in green algae, and its role in algal metabolism, remains limited and fragmented. This knowledge gap hinders the development of strategies for improving exoelectrogenesis-based biotechnologies.

We recently identified a green microalgal strain *Parachlorella kessleri* MACC-38, which exhibits unprecedented exoelectrogenic activity among green microalgal strains (Petrova et al., 2024). Under standard conditions (e. g. without iron limitation, which was previously used to promote exoelectrogenesis (Xue et al., 1998, Weger and Espie, 2000; Gonzalez-Aravena et al., 2018)), *P. kessleri* MACC-38 produced up to 10 times more electric current production than *C. reinhardtii* and the highly exoelectrogenic cyanobacterial species *Trichodesmium erythraeum*, and 3-4 times more current than other *P. kessleri* strains (Shlosberg et al., 2022; Petrova et al., 2024; Wroe et al., 2024).

The high exoelectrogenic capacity of *P. kessleri* MACC-38 suggests its potential not only for bioelectricity production but also as a model microorganism for studying the physiological roles of exoelectrogenesis and elucudating the metabolic processes that contribute to efficient photosynthetic exoelectrogenicity. In this study, we performed comparative photosynthetic characterization of MACC-38 and three less productive, *P. kessleri* strains (27.87, 211-11g and 211-11c). Furthermore, we investigated the effect of the exogenous electron acceptor FeCN on the photosynthetic performance of MACC-38. Our results suggest that a key distinctive feature of MACC-38, compared to less productive strains, is the crucial role of chlororespiration in balancing the redox state of the PETC and the intricate interplay between the PETC and the OPPP in providing NADPH as the direct electron donor for exoelectrogenesis.

## 2. Materials and methods

### 2.1. Algal cultures

Strain MACC-38, belonging to the *P. kessleri* species (Petrova et al., 2024), was acquired from the Mosonmagyaróvár Algal Culture Collection (Széchenyi István University, Hungary). *P. kessleri* strains 27.87, 211-11g and 211-11c were obtained from the Culture Collection of Algae at the University of Göttingen (SAG, Germany).

Algal suspension cultures were cultivated as described previously (Petrova et al., 2024). For all experimental procedures, with the exception of inductively coupled plasma mass spectrometry (ICP-MS) sample preparation, 3-day old cultures (in the early exponential growth phase) were harvested by centrifugation (1800×g, 5 min) and resuspended in Tris-phosphate medium (TP).

Where indicated, FeCN (1 or 5 mM) was added to cell suspensions at a density of 10 or 100 µg *Chl* (*a*+*b*)/ml, respectively. The suspensions were then incubated for 20 min at moderate light intensity of 150 µmol photons m^−2^ s^−1^. In experiments involving the inhibition of plastid terminal oxidase (PTOX), cell suspensions with Chl concentration 100 µg/ml were pre-treated with 2 mM propyl gallate (PG) for 20 min in the dark or at moderate light. Respective controls were incubated with 0.05% dimethyl sulfoxide (DMSO).

### 2.2. Chronoamperometric (CA) measurements

Electric current production by *P. kessleri* strains was measured for 10 min in the dark or at moderate light intensity of 150 μmol photons m^−2^ s^−1^ in the presence of 5 mM FeCN as an electron mediator, as described previously (Petrova et al., 2024).

### 2.3. Photosynthetic activity measurements

#### 2.3.1. Pulse-amplitude modulated (PAM) measurements

The steady state chlorophyll (Chl) *a* fluoescence signal and absorbance changes at ΔA_875-830 nm_, reflecting the redox state of P700 (the primary PSI electron donor), were monitored using a Dual-PAM-100 instrument (Heinz Walz, Germany). Except for experiments under anoxic conditions, all Chl *a* fluorescence and P700 measurements were performed with *P. kessleri* cells (40 μg *Chl* (*a*+*b*)) deposited onto Whatman GF/B glass microfibre paper after a 5-min dark adaptation. Anoxia was induced as per Ho et al. (2022) by adding 50 mM glucose, 10 U glucose oxidase and 30 U catalase to *P. kessleri* suspensions with Chl (*a*+*b*) concentration of 20 µg/ml. The suspensions were placed in a cuvette, covered with a layer of mineral oil and incubated in the dark for 20 min with continuous stirring.

Electron transport rate through PSII and PSI (ETR(II) and ETR(I), respectively), donor- and acceptor-side limitation of PSI (Y(ND) and Y(NA), respectively) were determined according to Baker (2008) and Klughammer and Schreiber (2008). Rapid light-response curves were constructed by plotting ETR(II) and ETR(I) against red actinic light (AL) intensities ranging from 25 to 1031 μmol photons m^−2^ s^−1^ (1-min exposure per light intensity).

The post-illumination Chl *a* fluorescence rise (PIFR) was monitored after 2 min of illumination with AL of 140 μmol photons m^−2^ s^−1^.

The kinetics of P700 oxidation was followed during application of AL with intensity 140 µmol photons m^−2^s^−1^ for 4 minutes

#### 2.3.2. Fast chlorophyll a fluorescence induction transients

Fast Chl *a* fluorescence (OJIP) transients were induced with a 3-second saturating light pulse (3500 μmol photons m^−2^ s^−1^, 625 nm peak wavelength) and recorded using a Handy-PEA (Hansatech Instruments Ltd., UK).

#### 2.3.3. Modulated reflection at 820 nm

The initial oxidation and reduction of PSI reaction centers was followed with M-PEA instrument detecting the modulated reflection (MR) at 820 nm. The probe light was provided by pulse-modulated LED source peaking at 820±25 nm, with 100% intensity. The MR signal was presented relative to the MR value recorded at the onset of illumination (MR_0_).

### 2.4. NAD(P)H fluorescence detection

NAD(P)H fluorescence (indicative of NAD(P)^+^ reduction) was monitored during dark-light transition using the NADPH/9-AA module for DUAL-PAM-100 (Heinz Walz, Germany) fluorometer (excitation: 365 nm, fluorescence detection: 420 - 580 nm). *P. kessleri* suspension cultures (10 µg Chl /ml) were dark-adapted for 5 min before the NAD(P)H fluorescence measurements. Light-induced NAD(P)^+^ reduction was observed under 140 µmol photons m^−2^s^−1^ red actinic light.

### 2.5. Immunodetection of photosynthetic proteins

Photosynthetic proteins were detected via Western blot analysis of membrane-enriched fraction isolated from 3-day old *P. kessleri* cultures. Algal suspensions (150 µg Chl) were pelleted by centrifugation at 12300×*g* and flash-frozen in liquid nitrogen with 30 µL of glass beads. All steps of the protein extraction protocol were performed at 4° C. Frozen pellets were thawed with 500 µL ice-cold extraction buffer (50 mM HEPES/KOH (pH 7.5), 10 mM sodium acetate, 5 mM magnesium acetate, 1 mM EDTA, 1 mM dithiothreitol, and 1×cOmplete protease inhibitor (Roche)). The samples were then vortexed for 2-3 min in 15-sec intervals. Next, the samples were shaken by a MM400 mixer mill (Retsch, Germany) at frequency of 300 Hz for 5 min. This step was followed by sonication for 30 sec at 60% amplitude with EpiSphear Probe Sonicator (120 W, 20 kHz, Active Motif, USA). The samples were centrifuged at 12000 rpm for 10 min. The resulting pellets were resuspended and vortexed in 350 µL of buffer containing 50 mM Tris-HCl, 0.28% Triton X-100, 1 mM dithiothreitol, and 1×cOmplete protease inhibitor. The samples were then shaken at 1100 rpm for 30 min, vortexed briefly and centrifuged at 15000 rpm for 10 min. The supernatant containing the membrane-enriched fraction was transferred to a clean tube. An amount of sample equivalent to 2.5 μg (100%), 1.25 μg (50%) and 0.625 μg (25%) *Chl* (*a*+*b*) was mixed with 6×Laemmli buffer (375 mM Tris-HCl (pH 6.8), 60% (v/v) glycerin, 12.6% (w/v) sodium dodecyl sulfate, 600 mM dithiothreitol, 0.09% (w/v) bromophenol blue) and denatured at 75 °C for 10 min prior to electrophoresis. The protein separation by SDS-PAGE electrophoresis and the wet transfer to a polyvinylidene difluoride membrane were previously described in Podmaniczki et al. (2021). Polyclonal primary antibodies raised in rabbit against PsaA, PsbA and PetB were purchased from Agrisera AB (Sweden). The incubation with the primary and the secondary antibodies (Bio-rad goat anti-rabbit IgG horseradish peroxidase conjugate) were performed according to the manufacturer’s instructions. Immunochemical detection of the proteins was done with SuperSignal West Pico PLUS Chemiluminescent Substrate (ThermoFisher Scientific, USA).

Densitometry analyses were performed with ImageJ software.

### 2.6. Determination of NADH/NAD^+^ and NADPH/NADP^+^ ratios

NADPH/NADP^+^ and NADH/NAD^+^ were quantified by the method of Gibon and Larher (1997) with modifications in the extraction procedure which include cell breaking in the presence of 1 M NaOH (for NADPH and NADH) or HCl (for NADP^+^ and NAD^+^) and an additional step of purification of the neutralized extracts by using Amicon Ultracel 10K centrifugal filter (Merck Millipore). The NADP(H) and NAD(H) concentrations were determined from the absorption at 570 nm with a calibration curve ranging from 10 to 100 pmol NAD(P)H.

### 2.7. Inductively-coupled plasma mass spectrometry

Prior to inductively-coupled plasma mass spectrometry (ICP-MS) measurements, algal suspensions were washed four times with ice-cold phosphate-buffered saline (PBS, pH 7.8), finally centrifuged at 1800×g for 5 min and cryogenically dried with Scan Speed 40 centrifugal evaporator and ScanVac CoolSafe freeze dryer (Labogene, Denmark). The dry weight of the samples was determined with an Entris 224 analytical balance (Sartorius, Göttingen, Germany). The dry algae samples were then digested for 2 hours in a mixture of 3 mL of concentrated (67 wt.%) HNO_3_ and 1.5 mL of concentrated (30 wt.%) H_2_O_2_ at 180°C.

ICP-MS measurements were carried out using an Agilent 7700x ICP-MS instrument (Agilent, Santa Clara, USA) equipped with an Agilent I-AS autosampler and a sample introduction system consisting of a MicroMist pneumatic nebulizer and a Peltier-cooled (4 °C) Scott-type spray chamber. Sample uptake rate and nebulizer gas flow rate was set to 750 mL·min^−1^, and 1.05 L·min^−1^, respectively. The plasma torch was operated with 1550 W RF forward power and 15 L·min^−1^ of Ar gas flow (plasma gas). All measurements were executed using Agilent MassHunter instrument control and data aquisition software (Agilent, Santa Clara, USA) which was set to analog mode, using an integration time of 1 second. The determination of Fe content of the algal samples were carried out by monitoring the signal of ^56^Fe isotope. To avoid spectral interferences, all measurements were performed using the Agilent ORS^3^ collision cell in He mode. ^45^Sc isotope was used as internal standard for Fe.

The ICP-MS was operated using high purity Ar (99.996%) and He (99.999%) gas obtained from Messer Hungarogáz Kft. (Messer Hungarogáz kft., Budapest, Hungary). Ultratrace-analytical grade nitric acid and hydrogen peroxyde (VWR, Radnor, USA) was used for the digestion of the samples. All samples were prepared using trace-analytical purity de-ionized water obtained from a MilliPore Eli× 10 device equipped with a Synergy polishing unit (Merck GmbH, Rahway, Germany). Standard solutions used for instrument calibration were made using certified stock solutions purchased from Inorganic Ventures Ltd. (Inorganic Ventures, Christiansburg, USA). For this purpose, the IV-ICPMS-71A multielement stock solution and the ICPMS-71D ICP-MS internal standard solution were used.

### 2.8. Membrane inlet mass spectrometry

Membrane inlet mass spectrometry (MIMS) measurements were performed using a MS-GAS-100 instrument (Photon Systems Instruments, Drásov, Czech Republic). The instrument is equipped with a 30 cm long polydimethylsiloxane (PDMS) inlet membrane probe consisting of a 1/8” tube, a Let-Lok reduction adapter (1/8”x1/16” OD), a 1/16” tube, a small push spring, a metal end piece (cap), and a PDMS membrane (approximately 3-4 cm long). The inlet membrane probe is connected to the instrument through a modular tubing consisting of 1/8” stainless tubes with 1/8”-1/8” UT adapters. The inlet membrane probe was submerged in a DW2/2 Clark-type liquid-phase oxygen electrode chamber (Hansatech Instruments Ltd, UK) with approximately 3.5 ml of cell suspension or culture medium. Stirring was provided with a magnetic stirrer (MAG1, Hansatech Instruments Ltd, UK). Light was provided through the side port of the oxygen electrode chamber using a KL 1500 Schott lamp. For liquid measurements, an integrated cooler water freezing trap, set to -80 °C, was used to minimize water vapour in the measuring system and protect the mass analyser.

Calibration of the instrument was performed according to the manufacturer’s standard operation procedures, before starting the measurements. This involved a background determination (at the internal inlet temperature of -80 °C) with the inlet valve closed to determine the ion currents of residual gases in the analysis chamber. These background ion currents were substracted from subsequent sample measurement. Subsequently, ion source tuning, mass scale tuning and adjustment, and offset calibrations were performed in TP medium (equilibrated with ambient air) in order to optimize ion source performance, determine the exact position of the peak maximum, and compensate for any drift of the measuring amplifier, respectively. Gas-specific calibration to convert ion currents into gas concentrations was performed using TP medium equilibrated with standard ambient air (400 ppm CO_2_).

N_2_ (m/z=28), O_2_ (m/z=32), Ar (m/z=40), and CO_2_ (m/z=44) were measured using the Faraday detector in both multiple ion detection (MID) and multiple concentration detection (MCD) modes to simultaneously record changes in ion currents and calibrated concentrations. Total pressure was also continuously monitored. Changes in O_2_ and CO_2_ concentrations were normalized to Ar to correct for signal drifts.

*P. kessleri* suspensions were set to a chl(*a*+*b*) concentration of 10 µg/ml. In the first 10 min of the measurements, the suspensions were kept in the dark. Subsequently, the MIMS signal was detected for 20 min under moderate light conditions (140 μmol photons m^−2^ s^−1^). For FeCN treatment, the cells were incubated with 1 mM FeCN for 10 min in the dark before the onset of the MIMS experiments.

### 2.9. Glucose 6-phosphate dehydrogenase (G6PDH) activity

To determine G6PDH activity, we prepared crude cell extracts from *P. kessleri* suspensions as in Zheng et al. (2017), immediately after 20-min dark incubation or after further exposing the cells to moderate light for 10 min. The suspensions used for crude extract preparation had cell density equivalent of 100 µg Chl (*a*+*b*)/ml. To assess the effect of FeCN, it was added to a concentration of 5 mM before the dark adaptation. The ability of G6PDH contained in crude cell extracts to reduce NADP^+^ was determined according to Zheng et al. (2017). The rate of NADP^+^ reduction was normalized to the total protein in the crude extracts as evaluated by the Bradford microassay (Kruger, 2002).

### 2.10. Statistical analysis

All data is presented as mean values of at least three biological replicates. Statistically significant differences to the respective controls were established in OriginPro 9.0 Software applying one-way ANOVA with Holm-Sidak post-hoc test at p < 0.05.

## Results

### 3.1. Electric current production by four P. kessleri strains

The exoelectrogenic activity of *P. kessleri* strains MACC-38, 211-11g, 211-11c and 27.87 was assessed using chronoamperometry (CA) (Fig. 1). Consistent with our previous findings (Petrova et al., 2024), within 10 min in the light, strain MACC-38 demonstrated the highest FeCN-mediated total current density production - 167 µA cm^−2^ mg^−1^ Chl, several times greater than previously reported for *C. reinhardtii* grown under non-stress conditions (Anderson et al., 2016). Strains 211-11g and 211-11c showed 3 times lower current densities (around 30 µA cm^− 2^ mg^−1^ Chl). Strain 27.87 showed intermediate productivity (approximately 70 µA cm^−2^ mg^−1^ Chl). Dark current densities ranged from 30-70 µA cm^−2^ mg^−1^ Chl for strains 27.87, 211-11g, and 211-11c, and reached approximately 124 µA cm^−2^ mg^−1^ Chl for MACC-38. Light-induced current (calculated as the arithmetic difference between total current obtained in the light, and the dark current) was approximately 55 µA cm^−2^ mg^−1^ Chl for MACC-38 and 10-12 µA cm^−2^ mg^−1^ Chl for the three other strains (Fig. 1). These results demonstrate significant variation in exoelectrogenic potential among *P. kessleri* strains, highlighting MACC-38 as a particularly promising candidate for bioelectrochemical applications.

**Fig. 1.**
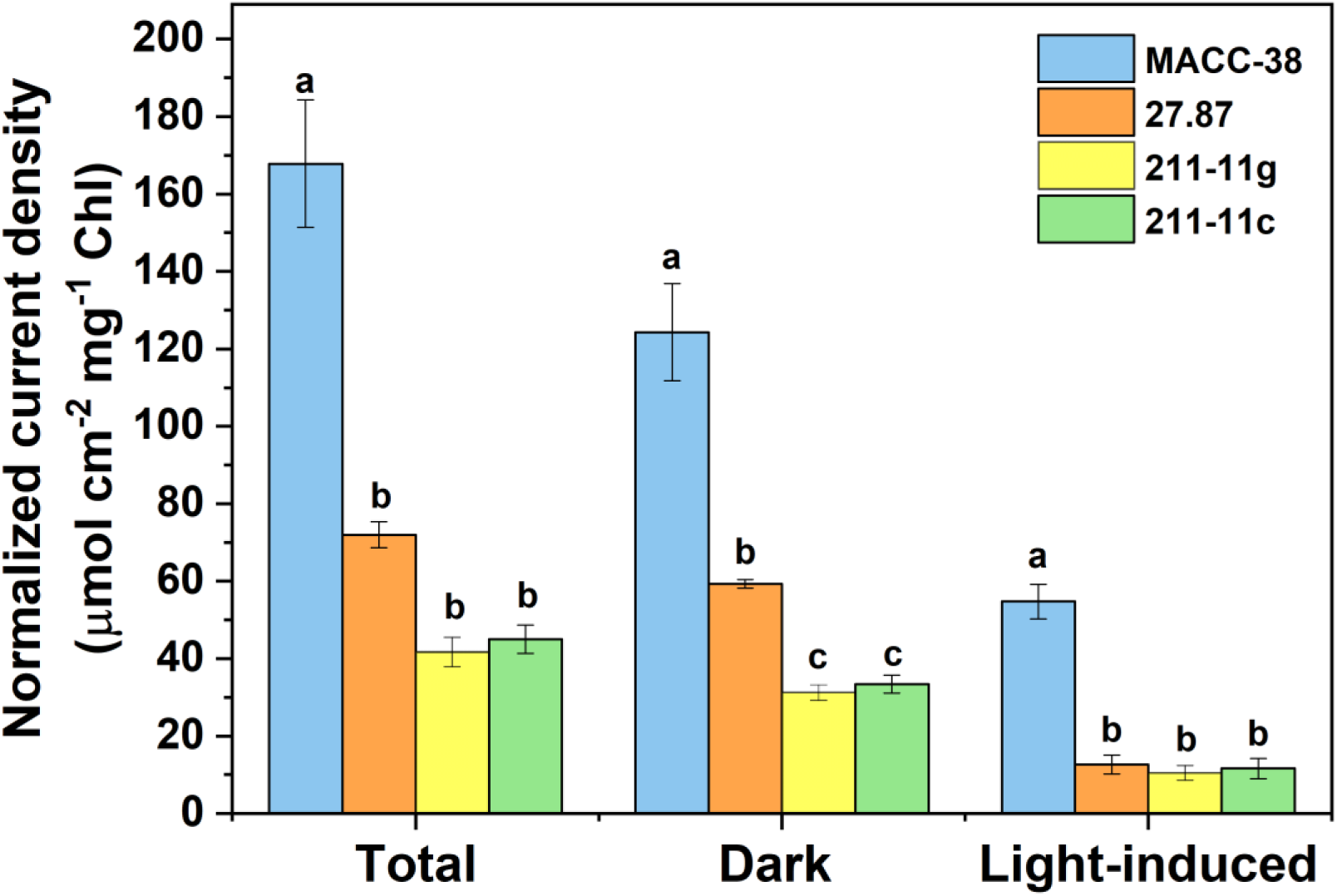
Electric current density generated by four *P. kessleri* strains normalized to chl(*a*+*b*) concentration. The electric current was recorded for 10 min in the dark and at moderate light (150 µmol m^−2^ mg^−1^ Chl). The current obtained in the light is referred to as total current and the light-induced current is the arithmetic difference between total and dark current. Statistically significant difference among the strains is denoted by distinct letters. Average values ± SEM, n=3.

### 3.2. Donor-side limitation of PSI in the MACC-38 P. kessleri strain

Given the pronounced, light-dependent exoelectrogenic activity of MACC-38 observed in our previous work (Petrova et al., 2024) and the data presented here (Fig. 1), we investigated whether this strain possesses unique photosynthetic electron transport characteristics. To characterize the photosynthetic apparatus of *P. kessleri* strains exhibiting varying exoelectrogenic activity, we measured electron transport rates of photosystem I (PSI) and photosystem II (PSII) across a range of light intensities (25-1000 μmol photons m^−2^ s^−1^, Fig. 2A, B).

**Fig. 2.**
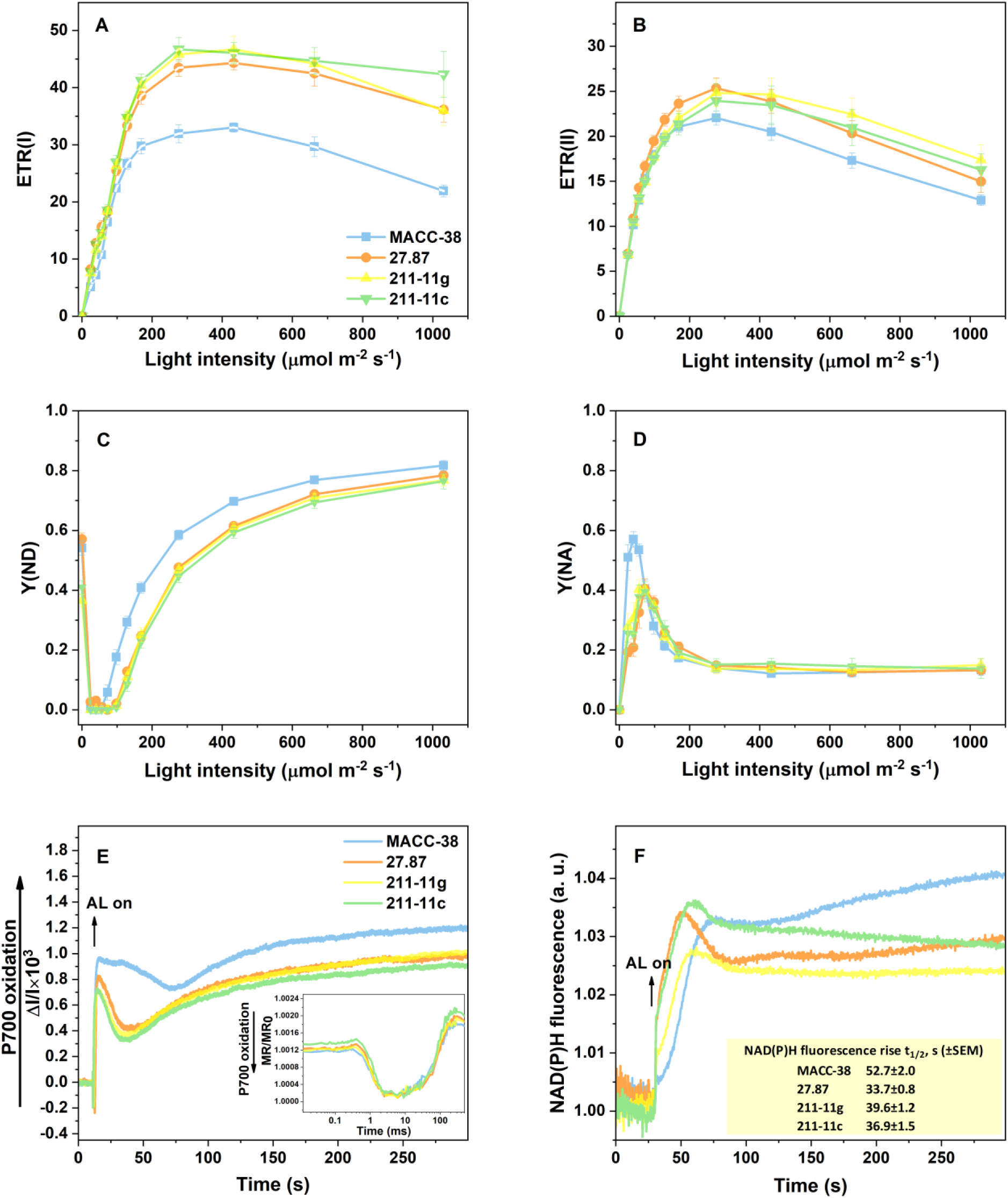
Photosynthetic electron transport characteristics of four *P. kessleri* strains. ***A***, Electron transport rate of PSI. ***B***, Electron transport rate of PSII. ***C***, Donor-side limitation of PSI. ***D***, Acceptor-side limitation of PSI. ***E***, Redox state of PSI during illumination with moderate light (150 µmol photons m^−2^ s^−1^), inset: representative curves of the initial oxidation and reduction of PSI shown on a logarithmic timescale. ***F***, Representative curves of NAD(P)H fluorescence kinetics during dark-light transition; inset: halftimes (t_1/2_) of NAD(P)H fluorescence rise in the light. Data in panels A-F and the inset of F represent averages ± SEM of at least 4 independent biological replicates.

MACC-38 had lower PSI electron transport rate ETR(I) compared to strains 27.87, 211-11g, and 211-11c (Fig. 2A). This difference became substantial above 130 μmol photons m^−2^ s^− 1^, a range of particular relevance as light-induced current production was measured at 150 μmol photons m^−2^ s^−1^ (Petrova et al., 2024). All *P. kessleri* strains showed similar ETR(I) kinetics: a rapid increase up to approximately 100 μmol photons m^−2^ s^−1^, followed by slower rise to a maximum around 300 μmol photons m^−2^ s^−1^, and then a decline at higher light intensities (Fig. 2A).

All *P. kessleri* strains showed similar ETR(II) curves in both shape and amplitude (Fig. 2B), reaching a maximum at approximately 300 μmol photons m^−2^ s^−1^. Beyond this point, MACC-38 exhibited a slightly lower ETR(II) than the other strains (Fig. 2B). However, this difference was not statistically significant.

To further elaborate on the electron transport through PSI, we examined donor-side (Y(ND)) and acceptor-side (Y(NA)) limitations (Fig. 2C, D). At light intensities above 70 μmol photons m^−2^ s^−1^, MACC-38 demonstrated markedly higher Y(ND) values than strains 27.87, 211-11g, and 211-11c (Fig. 2C). This difference was most pronounced between 100 and 170 μmol photons m^−2^ s^−1^. At light intensities above 50 μmol photons m^−2^ s^−1^, Y(NA) values were similar across all *P. kessleri* strains (Fig. 2D). Below 70 μmol photons m^−2^ s^−1^, however, MACC-38 exhibited substantially higher Y(NA) values than the other strains (Fig. 2D). These findings suggest that the reduced PSI electron transport observed in MACC-38 at moderate light intensities is primarily due to donor-side, rather than acceptor-side limitation.

To determine whether the differences in ETR(I) and Y(ND) are caused by variation in the rate of primary oxidation and reduction of PSI, we recorded the kinetics of modulated reflection (MR) at 820 nm for 500 ms (Fig 2E, inset). The absorbance changes at 820-830 nm primarily reflect changes in the P700 redox state, with contributions from PC and Fd (approximately 30 and 5%, respectively, Schansker et al., 2003). The slopes of the initial signal decrease at about 1 ms (reflecting the initial P700 oxiation, (Schansker et al., 2003, 2005; Strasser et al., 2010)) and the following rise starting at about 10 ms (corresponding to P700^+^ reduction by intersystem electron carriers (Schansker et al., 2003, 2005; Strasser et al., 2010)) were similar among the four *P. kessleri* strains (Fig 2E, inset). These results indicate that the relatively lower ETR(I) in MACC-38 is not caused by inherent difference in the rate of photochemical oxidation of PSI and in the inital re-reduction by the intersystem electron transporters occuring on a timescale of ms.

Furthermore, we examined the redox activity of PSI in continuous illumination by monitoring the absorbance changes at 830 nm for several minutes under actinic light of 150 μmol photons m^−2^ s^−1^ (Fig. 2E). In continuous light, after the initial oxidation of P700, the redox state of PSI is determined by simultaneously occuring re-reduction, re-oxidation, and gradual activation of electron sinks, giving the characteristic kinetics of the absorption signal at 830 nm (Harbinson and Hedley, 1993; Schansker et al., 2003; Belyaeva et al., 2019). Thus, the initial photoxidation and reduction of P700 were followed by a second phase of P700 oxidation (Fig. 2E), which was significantly larger in MACC-38 than in the other *P. kessleri* strains (Fig. 2E). Subsequently, P700^+^ became partially reduced by electrons originating from the intersystem electron transport chain (Harbinson and Hedley, 1993). The kinetics of P700^+^ reduction in MACC-38 demonstrate that this process occurred at a considerably lower rate in this strain than in the others (Fig. 2E). While in 27.87, 211-11g and 211-11c, the transient minimum of 830 nm change was registered at 35-40 s, in MACC-38 it occured at approximately 75 s. Further P700 oxidation, attributed to activation of the Calvin-Benson-Bassham cycle (CBB) (Harbinson and Hedley, 1993), was observed in all *P. kessleri* strains (Fig. 2E). The delayed reduction of P700^+^ by intersystem electron carriers is consistent with the higher donor-side limitation of PSI in MACC-38 compared to the other strains (Fig. 2C).

The lower PSI electron transport in MACC-38 may result in slower reduction of the terminal PSI electron acceptor, NADP^+^. To investigate the kinetics of photosynthetic NADP^+^ reduction, we examined NAD(P)H fluorescence transients in the four *P. kessleri* strains during a dark-light-dark transition (Fig. 2F). Upon illumination, *P. kessleri* strains 27.87, 211-11g and 211-11c exhibited similar NAD(P)H kinetics, with a fluorescence rise halftime of 33.7, 39.6 s and 36.9 s, respectively (Fig. 2F, inset). The NAD(P)H fluorescence kinetics of MACC-38 were substantially slower, with a halftime of approximately 52.7 s (Fig.2F, inset).

### 3.3. The donor-side limitation of PSI in MACC-38 is caused by lesser cytb_6_f abundance and increased chlororespiratory activity

Differences in photosynthetic complex stoichiometries could explain the enhanced PSI donor-side limitation in MACC-38. To investigate this possibility, we performed immunoblot analyses of PsbA, PsaA, and PetB, representing major subunits of PSII, PSI, and cytochrome b_6_f complexes, respectively (Fig 3A, B). We did not observe significant variations in PsbA and PsaA accumulation of the three *P. kessleri* strains relative to MACC-38. (Fig. 3A, B). However, PetB was up to twice less abundant in MACC-38 than in strains 211-11g and 211-11c. Strain27.87 had similar, but somewhat higher, PetB content than MACC-38 (Fig. 3A, B). Thus, the lower abundance of Cytb_6_*f* may contribute to the donor-side limitation of PSI in MACC-38, as observed using spectroscopy methods.

**Fig. 3.**
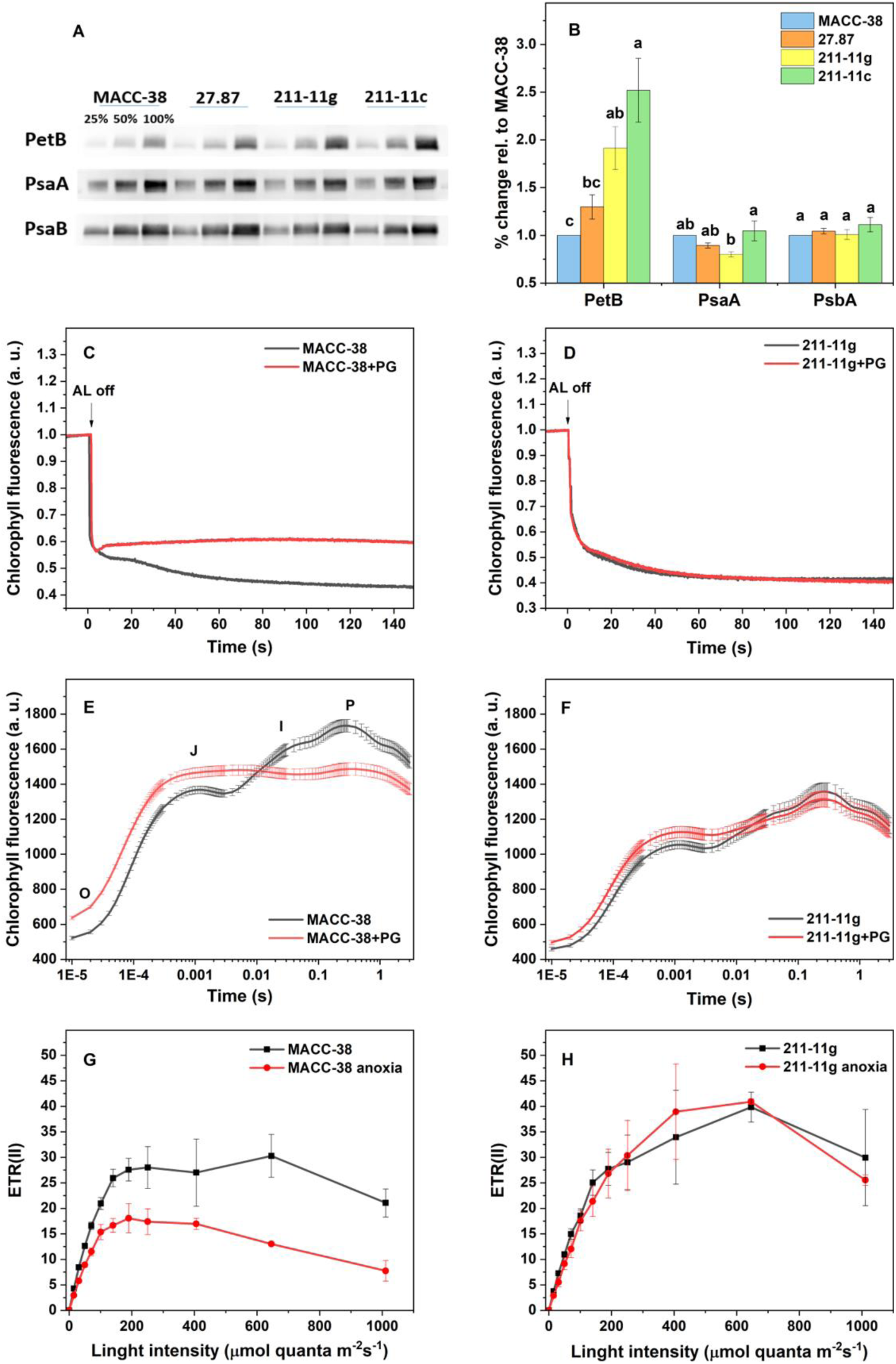
Analysis of PSI donor-side limitation in *P. kessleri* MACC-38. ***A***, Representative immunoblot images showing the abundance of photosynthetic subunits PetB, PsaA and PsbA in four *P. kessleri* strains. ***B***, Densitometry analysis of PetB, PsaA and PsbA immunoblots. Statistically significant differences among the strains are signified with different letters. ***C, D,*** Post-illumination fluorescence transients of MACC-38 (C) and 211-11g (D) after 20 min dark incubation, with 2 mM PG or 0.05% DMSO (control). ***E, F,*** Fast chl *a* fluorescence (OJIP) transients of MACC-38 (E) and 211-11g (F) after 20 min treatment with 2 mM PG or 0.05% DMSO (control) at moderate light (150 µmol photons m^−2^ s^−1^). ***G, H***, PSII electron transport rate in MACC-38 (G) and 211-11g (H) following anoxic treatment induced with glucose oxidase in the dark. ***B-H*** show average values ± SEM of at least 4 independent biological replicates.

When the intersystem electron transport, specifically the PQ pool, is in a reduced state, plastid terminal oxidase (PTOX) may mitigate this by acting as a plastoquinol:O_2_ oxidoreductase. This process is an alternative route for PQ oxidation, known as chlororespiration (Houille-Vernes et al., 2011, Houyoux et al., 2011). PQ oxidation by PTOX can be assessed by monitoring the post-illumination fluorescence rise (PIFR) in the presence of PTOX inhibitor propyl gallate (PG). The PIFR transients of all four *P. kessleri* strains showed similar features, with a fast initial drop of Chl *a* fluorescence, followed by a slow decrease (Fig. 1 SM), indicating two phases of oxidation of the PQ pool (Houyoux et al., 2011). The fast phase of PQ oxidation was, however, more pronounced in MACC-38 than the rest of the strains (Fig. 1 SM). Interestingly, upon PG treatment, the slow decline reflecting PQ-pool oxidation, was abolished in MACC-38, but not in 211-11g (for more clear presentation, here we compare MACC-38 only to 211-11g as a representative of the *P. kessleri* strains with lower exoelectrogenic activity Fig. 3C, D). This suggests remarkable PQ pool oxidation by PTOX in MACC-38, consistent with observations in tobacco and *C. reinhardtii* (Joet et al., 2002; Houyoux et al., 2011).

Fast Chl *a* fluorescence (OJIP) transients recorded following incubation of MACC-38 with PG under moderate light (150 μmol photons m^−2^ s^−1^ for 20 min) (Fig. 3E) showed increased F_0_ (O) values and a strong increase at the J step, approaching the maximum Chl fluorescence values (P or F_M_). These changes are characteristic of complete reduction of the PQ pool (Tóth et al., 2007), occuring in MACC-38 upon PG treatment in the light. Furthermore, the F_M_ value (Fig. 3E) and the F_V_/F_M_ parameter (an indicator of PSII activity), substantially decreased upon the PG treatment (Fig. 2 SM), suggesting the occurrence of photoinhibition upon PG treatment. These changes were minor in the other strains (for clarity, strain 211-11g is shown in Fig. 3F, 2 SM).

Since it has been reported that PG is not strictly specific to PTOX but can also inhibit the mitochondrial alternative oxidase (Cournac et al., 2000), we assessed dark respiration in the presence and absence of PG by membrane inlet mass spectrometry (MIMS). PG did not affect the respiration rate in the dark neither in MACC-38 nor in 211-11g, confirming that the observed effects of PG on Chl *a* fluorescence are due to its effect on the photosynthetic electron transport chain (Fig. 3 SM).

The role of O_2_ as electron acceptor in PQ-pool oxidation was further confirmed by anoxia treatment, in which cells were incubated with glucose, glucose oxidase, and catalase in the dark. A substantial decrease in ETR(II) was observed in anoxic MACC-38 cultures, but not in 211-11g (Fig. 3G, H). In summary, these results suggest that PTOX activity is remarkably high in the MACC-38 strain compared to the other strains, and this activity is essential to prevent over-saturation of the photosynthetic electron transport chain.

Furthermore, we performed chronoamperometric (CA) measurements of the four *P. kessleri* strains after pre-treatment with PG in the dark. In MACC-38, the total current in the light was 50% lower, the dark current was 35% lower, and the light-induced current decreased by 90% in the PG-treated cultures compared to the controls (Fig. 4A, B, C, respectively). In the other strains, the electric current density increased by up to 30% upon PG treatment, which was attributable to the increase in the dark current (Fig. 4A, B, C).

**Fig. 4.**
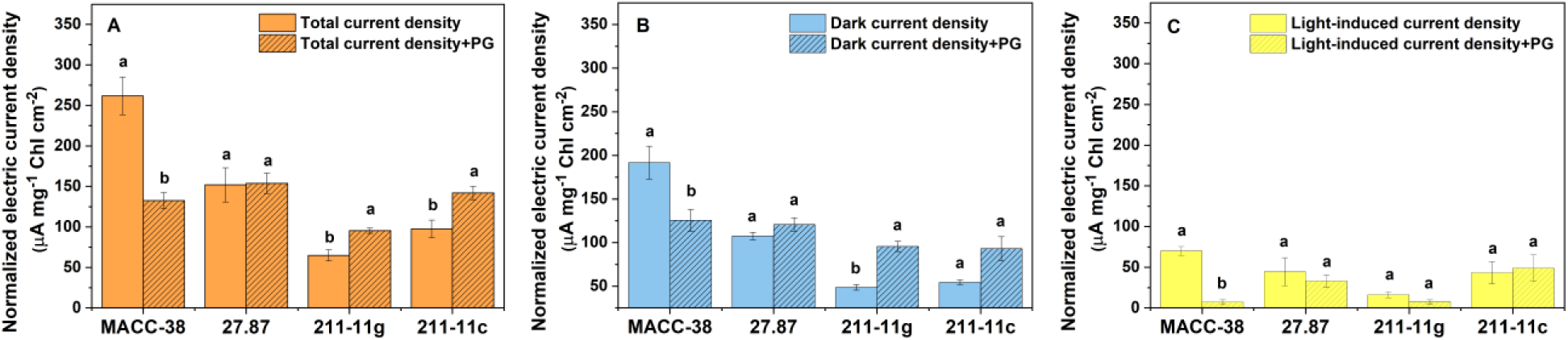
Effect of the PTOX inhibitor propyl gallate (PG) on the FeCN-mediated electric current production in four *P. Kessler* trains. ***A***, Electric current density under moderate light (150 µmol photons m^−2^s^−1^), normalized to chl(*a*+*b*) content. ***B***, Electric current density recorded in the dark. ***C***, Light-induced electric current density. Cells were pre-incubated in the dark for 20 min with 2 mM PG or 0.05% DMSO and washed twice with TP medium to remove the redox-active PG. Electric current was recorded for 10 min. Statistically significant difference between the PG-treated variants and the respective controls treated with 0.05% DMSO are denoted with different letters. Data in panels ***A-C*** show average values ± SEM of at least 4 independent biological replicates.

### 3.4. Extracellular FeCN reduction impacts photosynthetic electron transport and activates the oxidative pentose phosphate pathway

Due to the vast ability of FeCN to accept electrons from living algal cells (Gonzalez-Aravena et al., 2018; Petrova et al., 2024), FeCN treatment is an appropriate means of inducing electron export from the cells with the purpose to investigate the metabolic response to exoelectrogenesis. An important preliminary step, before studying the effects of FeCN on algal metabolism, is to establish whether it is involved in extracellular reactions only, or it can also enter green algal cells. It has been demonstrated that FeCN does not cross the cyanobacterial cell envelop (Kusama et al., 2021). However, an unequivocal proof that it only interacts with green microalgae at extracellular level is not available. Therefore, to find out whether FeCN can be internalized by *P. kessleri* cells, we measured their iron content after incubation with FeCN for 30 min or 2 h in the light, using inductively-coupled plasma mass spectrometry (ICP-MS). We did not observe any increase in cellular Fe content in either of the strains (Fig. 5A; for clarity, MACC-38 and 211-11g are shown), demonstrating that, at least on the time scale relevant for our experiments, FeCN is not internalized by *P. kessleri*. Therefore, the FeCN effects that we anticipate to observe are specifically due to its extracellular electron acceptor properties.

**Fig. 5.**
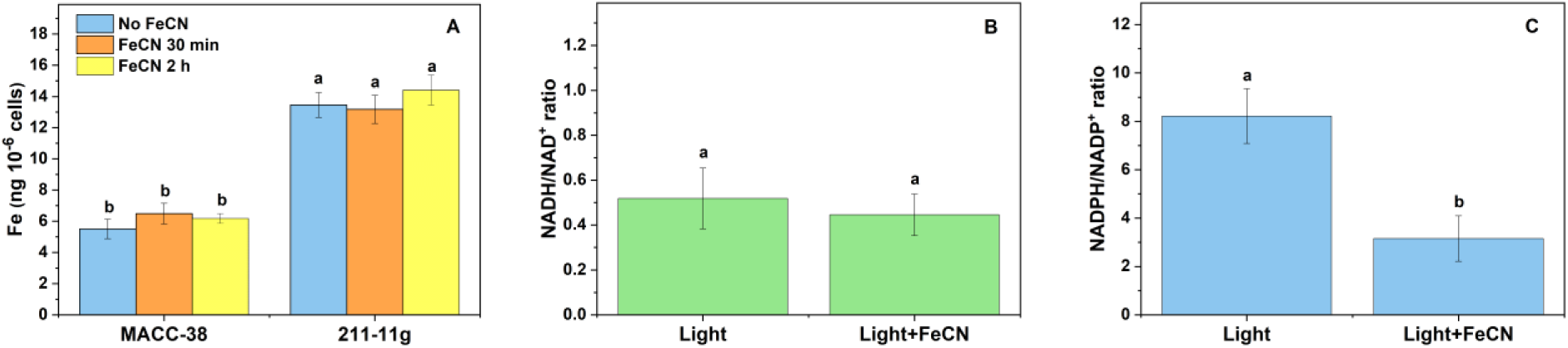
Effects of pre-incubation with FeCN in the light on *P. kessleri* strains MACC-38 and 211-11g strains. ***A***, Iron content of MACC-38 and 211-11g strains determined by ICP-MS before and after incubation with FeCN for 30 min and 2 h under moderate light (150 µmol photons m^− 2^s-^1^). ***B***, NADH/NAD^+^ ratio of MACC-38 after 20 min moderate light treatment with and without FeCN. ***C***, NADPH/NADP^+^ ratio of MACC-38 after 20 min moderate light treatment in the presence or absence of FeCN. Statistically significant differences between the FeCN-treated and untreated variants are denoted with different letters. Data in panels **A**-**C** show average values ± SEM of at least 4 independent biological replicates.

Previously described mechanisms of exoelectrogenesis in green microalga *C. reinhardtii* involve the oxidation of NADPH generated either at the acceptor side of PSI or through the OPPP, rather than NADH (Xue et al., 1998; Anderson et al., 2016). To determine the electron donor for exoelectrogenesis in MACC-38, we measured the NADH/NAD^+^ and NADPH/NADP^+^ ratios of this strain after treatment with AL (150 μmol photons m^−2^ s^−1^) in the presence or absence of FeCN (Fig. 5B, C). Light treatment in the presence of FeCN did not affect the NADH/NAD^+^ ratio significantly (Fig. 5B). However, adding FeCN led to a marked decrease in the NADPH/NADP^+^ ratio from 8.5 (light incubation without FeCN) to 3.2 (Fig. 5C). This results strongly indicate that the exoelectrogenic processes in MACC-38 show specificity for NADPH as a major electron donor over NADH.

We hypothesized that the substantial export of electrons and protons from MACC-38 cells during extracellular FeCN reduction (Petrova et al., 2024), along with the changes in NADPH/NADP^+^ ratio (Fig. 5C), could cause major alterations in the cellular redox status. To test this assumption, we investigated the effect of FeCN-pretreatment in the light on PETC. The chl *a* fluorescence induction transients of MACC-38 cultures treated with FeCN showed a slight increase at the J step (Fig. 6A), indicative of a partially reduced PQ pool (Tóth et al., 2007). This was not observed in the less exoelectrogenic strain 211-11g (Fig. 6B).

**Fig. 6.**
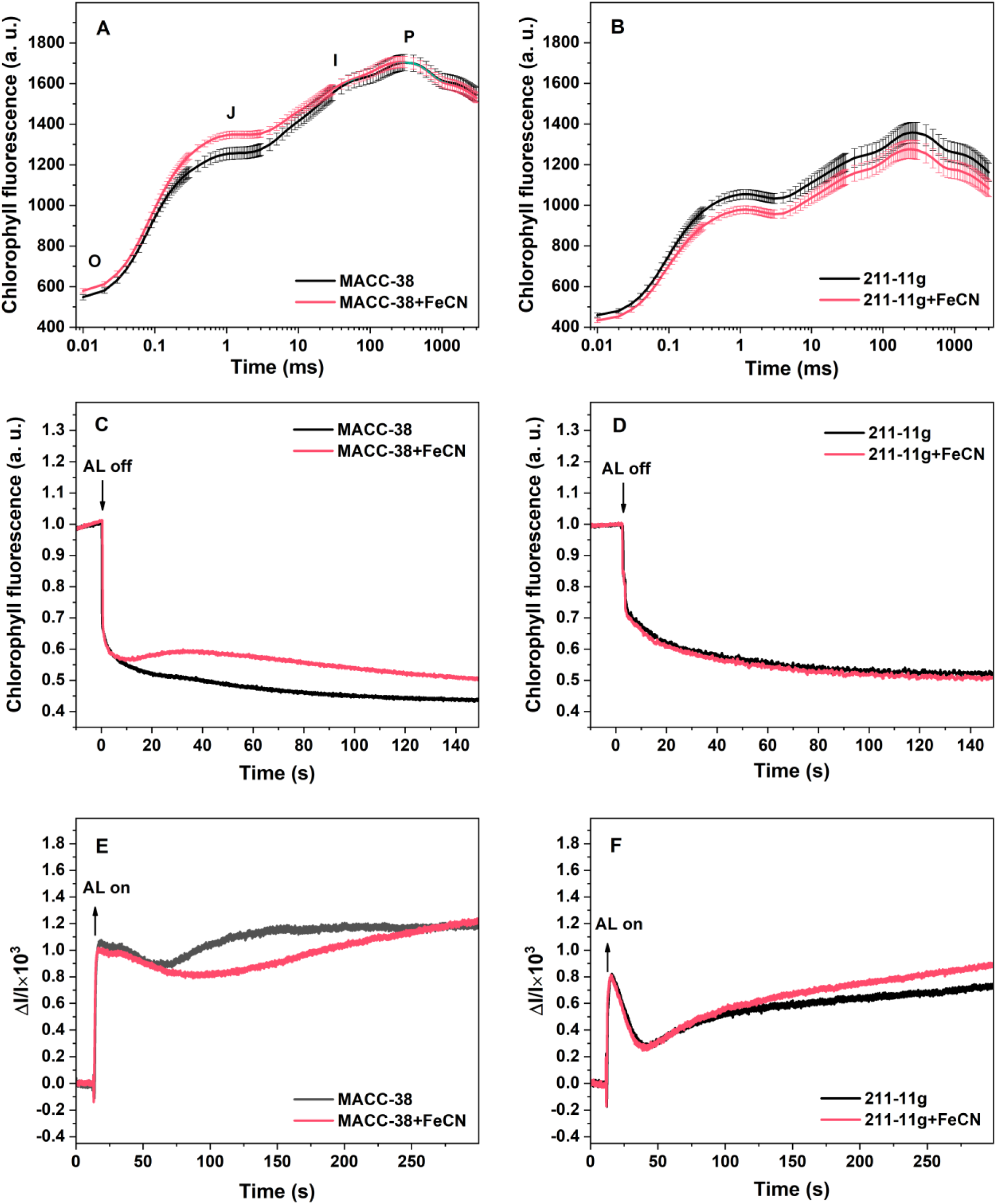
Effects of pre-incubation with FeCN for 20 min in the light (150 µmol photons m^−2^s^−1^) on the photosynthetic electron transport. ***A, B***, Fast chl *a* fluorescence (OJIP) transients of MACC-38 (A) and 211-11g (B). ***C, D***, Post-illumination fluorescence transients of MACC-38 (C) and 211-11g (D). ***E, F***, Kinetics of PSI oxidation/reduction (assessed by 820 nm absorbance measurements) in MACC-38 (E) and 211-11g (F). Data in panels **A**-**F** show average values ± SEM of at least 4 independent biological replicates.

The PIFR curves of FeCN-treated MACC-38 cells demonstrated a significant transient increase at about 40 s after switching off AL and an overall slow fluorescence decay (Fig. 6C), suggesting augmented PQ reduction (Houille-Vernes et al., 2011; Houyoux et al., 2011). On the other hand, the PIFR transient of FeCN-treated 211-11g cultures did not differ from the untreated suspensions (Fig. 6D).

Because NADP^+^ is the major electron acceptor of PSI, the increased oxidation of NADPH during FeCN reduction by MACC-38 may impact electron transport through PSI. To examine the effect of FeCN-mediated electron export on the PSI redox state, we monitored the absorbance changes at 830 nm under moderate AL after pre-incubation of MACC-38 with FeCN in the light (Fig. 6E). FeCN treatment did not affect the initial PSI oxidation and the rate of PSI re-reduction by the intersystem electron transport chain (observed as a dip at around 50-60 s after the onset of AL, Fig. 6E). However, PSI re-oxidation due to Calvin-Benson-Bassham (CBB) cycle activation, which occurs on the timescale of minutes (Harbinson and Hedley, 1993), was notably slower in FeCN-treated suspensions than in the control MACC-38 cultures (Fig. 6E). These effects of FeCN were not observed in the less exoelectrogenic strains (for clarity, 830 nm absorbance changes are shown only for the 211-11g strain in Fig. 6F).

The notable decrease in PSI re-oxidation rate in FeCN-treated MACC-38 cells prompted us to investigate whether extracellular FeCN reduction affects CO_2_ fixation, respiration, and O_2_ evolution. We used MIMS to detect the exchange of O_2_ and CO_2_ in the dark and at moderate light with or without FeCN (Fig. 7A, B). In spite of the slower P700 re-oxidation, FeCN did not inhibit CO_2_ fixation in MACC-38 in the light (Fig. 7B). Interestingly, although dark respiration was unaffected by FeCN (Fig. 7A), CO_2_ evolution was more than twice higher in FeCN-treated suspensions than in the control MACC-38 samples (Fig. 7B). In strain 211-11g, there was a statistically non-significant increase in CO_2_ evolution (Fig. 7B) with significantly higher respiration (Fig. 7A), suggesting that the reducing power for exoelectrogenesis in this strain may originate from respiration. In 211-11g suspensions, similar to MACC-38, FeCN did not significantly affect the rate of CO_2_ fixation, although photosynthetic O_2_ evolution showed a statistically non-significant increase (Fig. 7A, B).

**Fig. 7.**
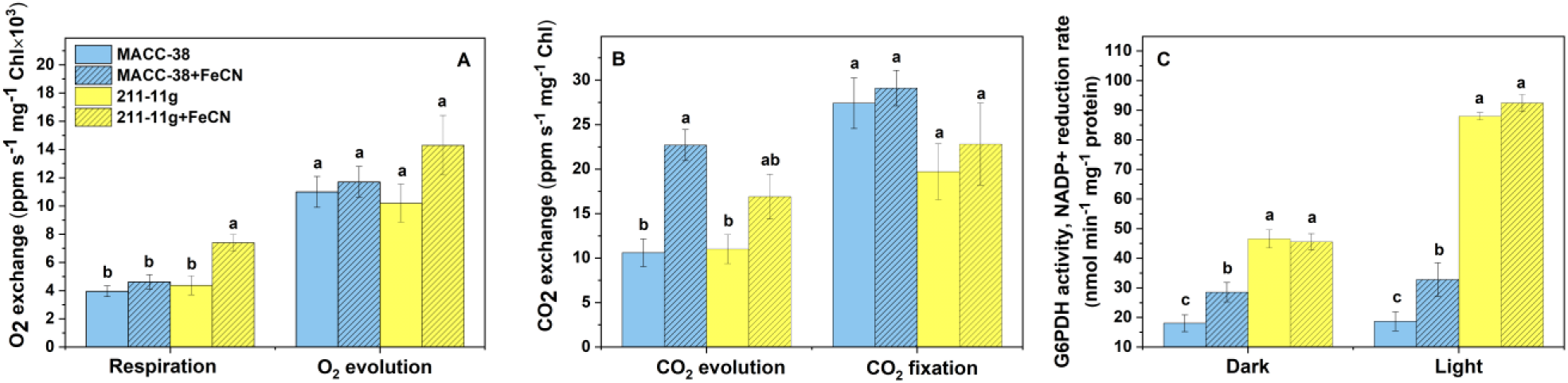
Effects of FeCN on O_2_ and CO_2_ exchange and G6PDH activity in *P. kessleri* strains MACC-38 and 211-11g. ***A***, Dark respiration and photosynthetic O_2_ evolution rates, measured by MIMS, with or without FeCN at moderate light (150 µmol photons m^−2^s^−1^). ***B***, Rates of dark CO_2_ evolution and CO_2_ fixation in the light (150 µmol photons m^−2^s^−1^), measured by MIMS with or without FeCN. ***C***, G6PDH activity in crude extracts of MACC-38 and 211-11g pre-incubated with FeCN in the dark and under moderate light (150 µmol photons m^−2^s^−1^). In **A** and **B** the O_2_ and CO_2_ % concentration signals were normalized to Ar signal and to chl(*a*+*b*) concentration. In **C** NADP^+^ reduction rates were determined between the 5 and 10th min of incubation of crude extracts with glucose 6-P and NADP^+^ at room temperature. Statistically significant differences between the two strains and the FeCN-treated and untreated variants are denoted with different letters. Data in panels **A**-**C** show average values ± SEM of at least 4 independent biological replicates.

Our results, suggesting that the exoelectrogenic activity in MACC-38 consumes reducing power specifically in the form of NADPH, and is accompanied by increased CO_2_ production, led us to focus on OPPP. This pathway, a major source of NADPH in plant tissues, is regulated by changes in the cellular redox state and NADPH/NADP^+^ ratio, and includes steps of CO_2_ release (Kruger and von Schaewen, 2003). Furthermore, OPPP activity is of paticular interest in studies on the light-induced current production by green microalgae because it has been demonstrated that, in *C. reinhardtii*, OPPP reactions take place predominantly in the chloroplast (Klein, 1986).

We quantified the activity of glucose 6-phosphate dehydrogenase (G6PDH), the first and rate-limiting step of the OPPP, in crude extracts from MACC-38 and 211-11g treated with FeCN under the same conditions as the MIMS measurements (dark followed by light incubation with or without FeCN). Incubation of MACC-38 with FeCN in the dark and in the light caused a 50% and a 100% increase in G6PDH activity, respectively, compared to untreated suspensions (Fig. 7C). In contrast, although the G6PDH activity of 211-11g was substantially higher than that of MACC-38, it remained unchanged regardless of FeCN treatment (Fig. 7C) supporting the hypothesis that respiration is a primary source of reducing power for exoelectrogenesis in this strain. Although these results do not preclude the possibility of other enzymes contributing to the increased CO_2_ release and NADP^+^ reduction in MACC-38 during FeCN reduction, they demonstrate that the availability of an external electron sink, such as FeCN, leads to metabolic changes in the cell, specifically, OPPP activation.

## 4. Discussion

Electron export in green microalgae has previously been linked to two processes: (i) extracellular superoxide production by *C. reinhardtii* under replete conditions (Anderson et al., 2016) and (ii) reduction of trace metals (Cu(II) and Fe(III)) upon iron and copper limitation in *Chlorella kessleri* (recently renamed to *Parachlorella kessleri*) and *C. reinhardtii* (Hill et al., 1996; Lynnes et al., 1998; Xue et al., 1998; Weger et al., 1999; Weger et al., 2009; Gonzalez-Aravena et al., 2018). Both superoxide generation and Fe(III)-reduction are upregulated by light and utilize NADPH as an electron donor (Xue et al., 1998; Lynnes et al., 1998; Anderson et al., 2016), implying the involvement of photosynthesis in exoelectrogenesis. Under iron-limiting conditions, the OPPP has been identified as another NADPH source for electron export (Xue et al., 1998). However, in general, the bioenergetics of exoelectrogenesis in green microalgae remains a largely unexplored field.

Recently, we identified a green microalgal strain, *P. kessleri* MACC-38, capable of producing, under replete conditions, 10 times higher electric current than *C. reinhardtii* and 3-4 times larger current than other *P. kessleri* strains (Petrova et al., 2024, Fig. 1). Although we could not definitively attribute the exoelectrogenic capacity of this strain to any of the known electron export processes, we found that approximately 70% of the light-induced current in MACC-38 is dependent on photosynthesis (Petrova et al., 2024). In this study, we characterize the photosynthetic electron transport in MACC-38 and three other, less productive *P. kessleri* strains to identify distinctive photosynthetic features associated with augmented exoelectrogenesis.

We found that in the highly electrogenic MACC-38 strain, NADPH fuels electrogenesis, similar to *C. reinhardtii* (Xue et al., 1998; Anderson et al., 2016, Fig. 5C). However, unlike less productive *P. kessleri* strains, MACC-38 exhibits significantly lower PSI electron transport (Fig. 2A) resulting in slower initial rate of NAD(P)^+^ reduction (Fig. 2F).

One reason for the donor-side limitation of PSI is the lower Cytb_6_f content in MACC-38 (Fig. 3A, B). However, the redox state of the intersystem electron transport depends not only on photochemical but also on non-photochemical redox reactions, including cyclic electron flow, carbohydrate catabolism (Jans et al., 2008; Johnson et al. 2014), and chlororespiration via PTOX (Houille-Vernes et al., 2011). We found that inhibition of PTOX by PG dramatically increases PQ-pool reduction in MACC-38 and anoxic conditions strongly inhibit ETR(II) (Fig. 3C, E, G), indicating that PTOX substantially oxidizes the PQ pool. This is particularly relevant given the low Cytb_6_f content (Fig. 3A, B). Unfortunately, we were not able to quantify PTOX in *P. kessleri,* neither with *C. reinhardtii* PTOX2, nor with *A. thaliana* PTOX antibodies. Nonetheless, our results suggest that chlororespiration plays a crucial role in oxidizing the PQ pool and contributes to the enhanced PSI donor-side limitation in the highly exoelectrogenous MACC-38 strain. Previously, PSI donor-side limitation has been linked to PTOX activity in *Marchantia polymorpha* (Messant et al., 2023).

Interestingly, inhibition of PTOX significantly decreased light-indiced electric current production specifically in MACC-38 (Fig. 4A, B, C). This decrease may result from the lower ETR(II) and increased reduction state of PQ pool observed upon PTOX inhibition (Fig. 3E, F), potentially inducing photosynthetic control over linear electron transport. However, a direct connection between exoelectrogenesis and chlororespiration in MACC-38 warrants further investigation. The absence of effect of PG on light-induced current in 27.87, 211-11g and 211-11c corroborates the less significant role of chlororespiration in these strains (Fig. 3D, F, H). The increased dark current observed in these strains after pre-treatment with PG may be explained by its potent antioxidant properties, which could have diverse effects unrelated to respiratory reactions (Medina et al., 2013).

FeCN is a commonly used electron mediator in BPV research (Anderson et al., 2016; Shlosberg et al., 2021, Wang et al., 2021; Petrova et al., 2024). Furthermore, as a potent electron acceptor, FeCN can be utilized to study the role of exoelectrogenesis in algal metabolism. Until now, it was uncertain whether, in addition to its function as extracellular electron acceptor, FeCN might also exert other effects by entering green algal cells. We demonstrated that FeCN is not internalized by *P. kessleri* by showing that the cellular iron content remains unchanged after FeCN treatment (Fig. 5A). Thus, the effects of FeCN on *P. kessleri* metabolism are solely due to its extracellular redox activity and its role to act as a potent electron acceptor. In agreement with previous reports, we show that exoelectrogenesis is linked to NADPH oxidation and increased NADP^+^ availability. Since NADP^+^ is the terminal acceptor of PSI, it could be expected that FeCN treatment would cause faster PSI re-oxidation. However, conversely, we observed that FeCN-treated MACC-38 exhibited a transient slowdown in PSI re-oxidation (Fig. 6E). We initially hypothesized that this slower PSI re-oxidation might result from competition between exoelectrogenic reactions and the CBB cycle for NADPH. However, this possibility was ruled out because CO_2_ assimilation in the light was unaffected by FeCN (Fig. 7B). Furthermore, we observed a significant delay in PQ-pool re-oxidation in the dark (Fig. 6C) and a slight over-reduction of the PQ pool upon application of a saturating light pulse (Fig 6B), which is consistent with the slower re-oxidation of PSI after FeCN treatment. Previously, extracellular reduction of FeCN was shown to serve as an electron sink, alleviating over-reduction of PETC in *C. reinhardtii* and *Chlorella pyrenoidosa* (Xue et al., 1998; Wang et al., 2021). However, our data points that, in the highly exoelectrogenic strain MACC-38, FeCN-reduction causes transient over-reduction of the photosynthetic electron transport.

The observed doubling of CO_2_ evolution in the dark in the presence of FeCN, without a corresponding change in respiration (Fig. 7A, B), aligns with the activation of the OPPP by electron export in MACC-38 cultures. Consistently, we found that FeCN increased the activity of G6PDH, rate-limiting enzyme of the OPPP, by approximately 50% in the dark and 100% in the light (Fig. 7C). As the OPPP in the green alga *C. reinhardtii* predominantly functions in the chloroplasts (Klein et al., 1986), and two of the reactions in this pathway involve NADP^+^reduction (Kruger and von Schaewen, 2003), we suggest that the transient slowdown of PSI re-oxidation may occur due to temporal depletion of the PSI end acceptor, NADP^+^, by the OPPP, or increased reduction of the PQ pool by NADPH (generated by the OPPP) during cyclic electron flow (CEF) around PSI. Moreover, previous studies have shown that increased G6PDH activity during salt stress in *Physcomitrella patens* is associated with increased CEF (Gao et al., 2015). These data demonstrate that a strong extracellular electron sink can significantly modify and regulate cellular metabolism. This has implications for both BPV research and our understanding of the physiological relevance of exoelectrogenesis.

G6PDH isoforms and their biochemical characteristics vary significantly among green microalgae. In *Chlorella sorokiniana* and *Koliella antarctica* a single G6PDH isoform is present, displaying similarity to the plastidic isoform in higher plants. These isoforms exhibit reduced sensitivity to NADPH inhibition and maintain significant activity under illumination, while being subject to redox regulation, via e.g., thioredoxin (Esposito et al., 2006; Ferrara et al., 2013). Conversely, *Chlorella vulgaris* possesses two distinct isoforms, resembling the cytosolic and chloroplastic isoforms found in higher plants. The cytosolic isoform operates independently of the cellular redox state, whereas the chloroplastic isoform is strongly inhibited in reducing conditions (Honjoh et al., 2006). Consequently, our finding of significant G6PDH activity under both dark and light conditions aligns with previous observations in closely related *Chlorella* species.

Green algae *C. reinhardtii* and *Selenastrum minutum* demonstrate an upregulation of OPPP, and specifically G6PDH, in response to conditions demanding increased reducing power. This is exemplified by NO_3_-assimilation in N-starved cultures, which drives a decrease in the NADPH/NADP^+^ ratio and subsequently activates the OPPP (Huppe et al. 1992, 1994; Esposito et al., 2006). Therefore, we hypothesize that the lowered NADPH/NADP^+^ ratio during FeCN-reduction triggers G6PDH (and OPPP) activation, thereby providing MACC-38 cells with the necessary reducing power. Interestingly, a similar effect was observed in *Nicotiana* species as a reaction to elicitors cryptogein and INF1, provoking extracellular superoxide production (and thus electron export), NADPH oxidation and OPPP activation (Pugin et al., 1997; Asai et al., 2011).

However, while increased electron export has been associated with OPPP activation in iron-limited *C. reinhardtii*, the metabolic state of iron-deprived cells differs profoundly from that of iron-replete cultures. Iron limitation induces substantial stress on the photosynthetic apparatus mostly due to reduced abundance of key iron-containing photosynthetic subunits, altering electron transport (Terauchi et al., 2010; Yadavalli et al., 2012; Devadasu et al., 2016). Therefore, iron-limited algae likely rely more heavily on non-photosynthetic pathways for reducing power, especially in conditions leading to increased NADPH oxidation. This is supported by the observation that, unlike *P. kessleri* grown under standard conditions, iron-limited *C. reinhardtii* exhibited near-complete inhibition of photosynthetic CO_2_ fixation during FeCN-reduction at subsaturating light (Weger and Espie et al., 2000). Furthermore, FeCN reduction did not provoke changes in CO_2_ and O_2_ exchange in iron-replete *C. reinhardtii* (Weger and Espie et al., 2000).

Although FeCN-reduction did not inhibit CO_2_ fixation or O_2_ evolution in MACC-38 under illumination (Fig 7A, B), we observed a transient over-reduction of PETC and a slowed re-oxidation of PSI due to acceptor side-limitation following FeCN treatment (Fig. 6A, C, E). These observations led us to propose a model of the metabolic reactions occurring during exoelectrogenesis, which explains the trainsient over-saturation of PETC with activation of OPPP caused by alterations in the cellular redox balance resulting from electron and proton export. OPPP activation may temporarily deplete NADP^+^, the terminal acceptor of PSI, or induce CEF, thus contributing to the slower re-oxidation of PSI in MACC-38. We also propose a potential role for chlororespiration in exoelectrogenesis in MACC-38. Oxidation of the PQ pool by PTOX may mitigate the over-reduction of PETC upon OPPP activation. However, further investigation is required to confirm the involvement of chlororespiration in exoelectrogenesis.

In summary, we have elucidated key metabolic mechanisms driving unprecedented exoelectrogenic activity in *P. kessleri* MACC-38. Our findings reveal that chlororespiration plays a critical role in maintaining PETC redox balance, while OPPP activation provides essential reducing power, with both pathways intricately linked to efficient electron export. We further demonstrated that FeCN, acting as a potent external electron acceptor, induces specific metabolic shifts. These results contribute to a more complete understanding of electron export in green microalgae, identifying the PETC-OPPP interplay as a crucial target for enhancing biophotovoltaic and bio-reduction technologies. Moving forward, further research should focus on optimizing these interactions to maximize solar energy capture and conversion in microalgal systems, ultimately paving the way for sustainable bioelectrochemical and biocatalytic applications.

## Acknowledgements

The Authors are grateful to Drs. Petar H. Lambrev and Parveen Akhtar for the technical assistance with the M-PEA instrument and 820 nm measurements and to Dr. Roland Tengölics for his help with algal biomass lyophilization. We thank Dr. László Kovács for the helpful discussions.

