## Supplementary material for "Unprecedented exoelectrogenic activity in *Parachlorella kessleri* MACC-38: The interplay of photosynthetic electron transport and the oxidative pentose phosphate pathway"

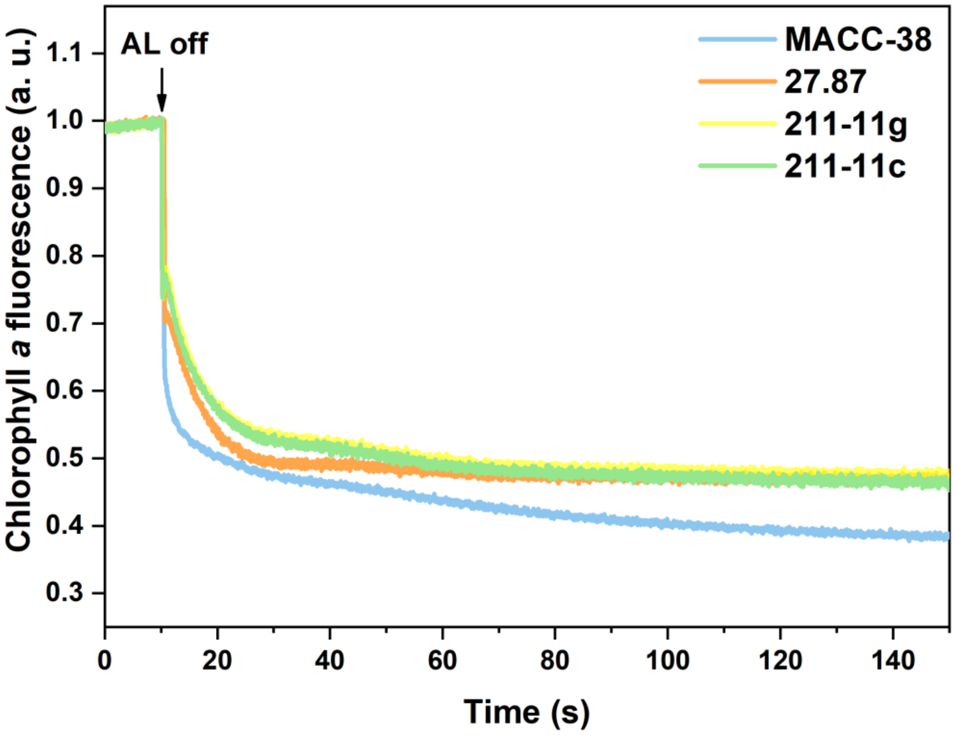


***Fig. 1 SM***. Post-illumination fluorescence rise (PIFR) of four *P. kessleri* strains exposed to 140 µmol quanta $m^{-2}$ $s^{-2}$ red actinic light for 2 min.

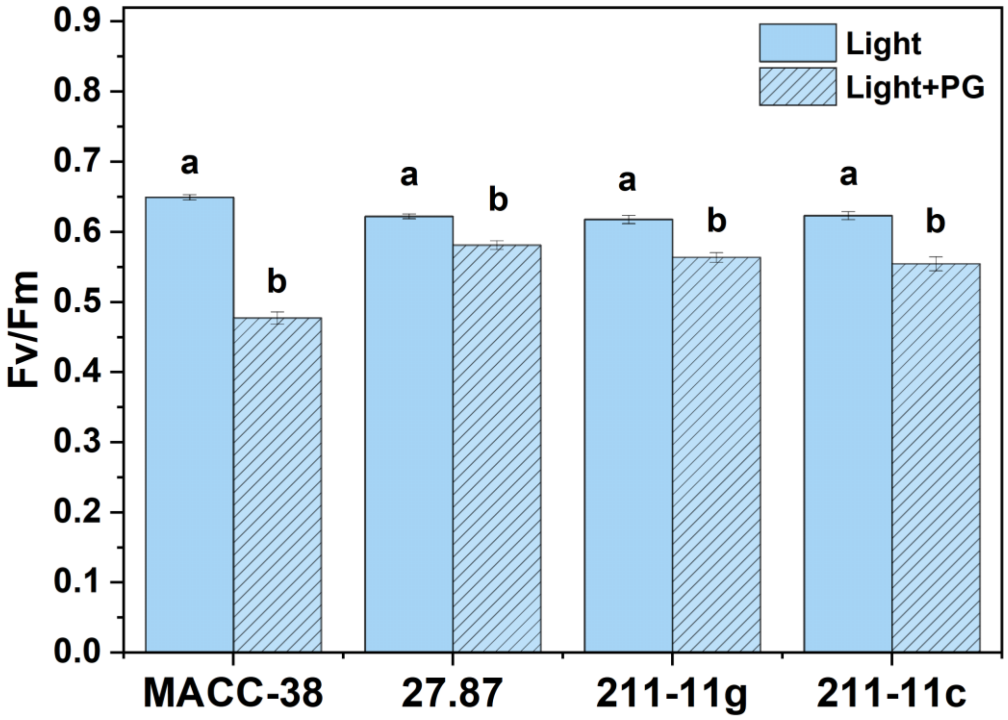


***Fig. 2 SM***. Maximum quantum efficiency of PSII ($F_V/F_M$) of four *P. kessleri* strains after treatment with PTOX inhibitor propyl gallate (PG) at moderate light (150 µmol quanta $m^{-2}$ $s^{-2}$). Controls were incubated with 0.05% DMSO.

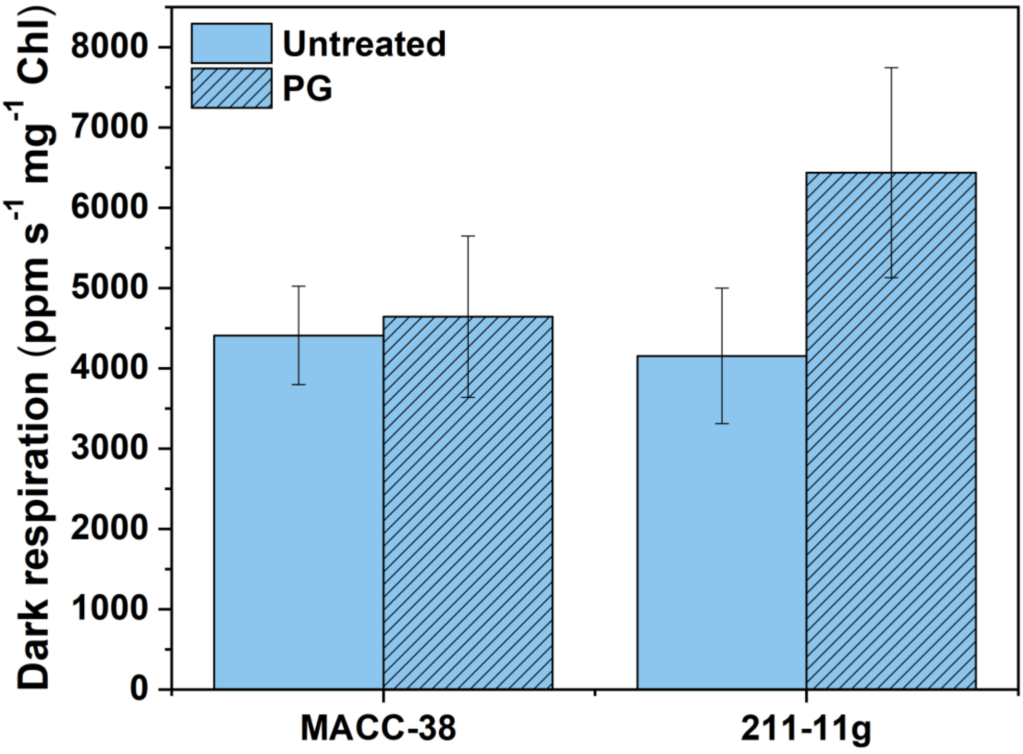


***Fig. 3 SM.*** Effect of PTOX inhibitor PG on the respiratory rate in the dark in *P. kessleri* strains MACC-38 and 211-11g detected with membrane inlet mass spectrometry (MIMS). PG was added to the algal suspensions (10 µg Chl (*a*+*b*)) at concentration 0.5 mM 10 min prior to beginning of MIMS measurement.
